# Collective membrane dynamics emerging from curvature-dependent spatial coupling

**DOI:** 10.1101/164392

**Authors:** Zhanghan Wu, Maohan Su, Cheesan Tong, Min Wu, Jian Liu

## Abstract

Membrane curvature has been recognized as an active participant of fundamental biological processes including vesicular transport and organelle biogenesis, but its effects on membrane remodeling are typically local. Here we show membrane curvature plays a critical role in propagating cortical waves and modulating mesoscale dynamics in living cells. We employ a membrane shape-dependent mechanochemical feedback model to account for the observed oscillatory travelling waves of Cdc42, F-BAR proteins and actin. We demonstrate that oscillatory membrane shape changes accompany and are required for such spatiotemporal patterns. In addition, modulating the curvature preference of the F-BAR proteins or membrane tension perturbs wave propagation. Our findings identify a distinct role of membrane curvature in mediating collective dynamics of cortical proteins and provide a molecular framework for integrating membrane mechanics and biochemical signaling in the context of subcellular pattern formation.

## Introduction

Dynamic patterns as waves are widely observed in many biological systems. They reflect the pulse of underlying signal transduction [1]. Actin waves on plasma membrane, which underlie many important cellular functions [2], are especially intriguing. Many different models have been proposed for waves of actin, most of which are based on reaction-diffusion mechanism [3,4] or its variants with additional factors including cytoplasmic flow [5], myosin-dependent actin transport [6,7] and motor-independent treadmilling [8,9]. Models with actin transport or treadmilling are likely not directly applicable to the actin waves observed in *Dictyostelium* [10,11] or immune cells [12], where sequential cycles of dendritic actin assembly and disassembly are involved [13]. These actin-centric wave models differ significantly in terms of whether actin provides both positive and negative feedbacks [14], positive feedbacks alone with an unknown diffusing inhibitor [15], or negative feedbacks alone with an upstream activator [4].

Commonly in these models, the plasma membrane is treated as a passive and flat two-dimensional manifold. However, it has been recognized that membrane shape change can feed back onto biochemical pathways in governing many cellular processes [16–20]. Interestingly, membrane curvature was theoretically predicted to induce traveling wave in active membrane when little was known about the molecular mechanisms of membrane curvature sensing or generation [21]. In this mechanical wave model, both types of curvature (concave and convex) are necessary and they act as positive and negative feedbacks, respectively. With the increasing interests in the involvement of actin waves in cellular processes involving membrane morphogenesis, such as cell migration, spreading and division, membrane geometry becomes an attractive parameter for modeling [22,23], as well as for experiments [24,25]. Membrane shape changes were also contemplated for actin wave-related phenomena such as cell edge protrusions [26,27] and dorsal ruffles, a wave-like propogation on the apical side of cell which involves membrane protrusions [28]. Because of a lack of knowledge of specific proteins for modulating membrane shape change, these models also differ in the specific requirements of mechanisms for curvature sensing or generation, where changes in membrane shape could be a passive result of forces applied by the actin filaments. More importantly, protrusion of the membrane by actin polymerization is an essential aspect of positive feedback and force generation mechanism. Therefore, it is not clear whether similar mechanisms could apply to actin waves propagating on the ventral surface of the cell, where membrane protrusion is presumably inhibited by the substrate. The role of membrane shape in actin waves at the basal surface, either passive or active, remains obscure due to a lack of direct experimental observation of plasma membrane deformation in these waves [25].

While all those models can generate morphologically similar wave patterns, they differ significantly in the mechanisms of wave propagation. In particular, whether actin plays an activating or antagonistic role, and whether membrane mechanics plays an active function in driving waves remain as fundamental open questions [14,29]. In this work, we combine theory and experiments to study the nature of collective membrane and protein waves in mast cells. Here we propose a ventral actin wave model in which the waves are driven by membrane shape undulation and fed back by curvature-dependent biochemical pathways, instead of driven by protrusive forces generated by actin polymerization. We show experimentally that membrane curvature is indispensable and that the sensitivity of cortical proteins to membrane shape renders the ventral cortex itself an excitable medium. Consequently, membrane shape deformation potentiates long-range traveling waves that emerges from this cortical excitability.

## Results

### Propagation Speed of Waves does not Scale with Actin Polymerization Rate

A number of actin wave models argue that protrusive force driven by actin polymerization on the membrane is the activating factor for travelling wave formation [14,26,29]. If the spatial propagation is driven by actin polymerization, wave propagation speed is expected to be proportional to the actin protrusion velocity, which depends on the rate of actin polymerization [2]. Our previous findings introduced a number of explicit components of the actin wave in mast cells [30]. Upon antigen stimulation, not only actin, but also a few membrane-interacting proteins, including N-WASP, Cdc42 and F-BAR domain-containing protein FBP17, exhibit coordinated traveling wave behavior within the ventral cortex. These proteins allow us to investigate the effect of perturbing actin turnover in cortical waves using reporters other than actin itself. With low doses of Latrunculin A (LatA), which prevents actin polymerization by sequestering G-actin, we observed that in mast cells, the oscillation period increased, but FBP17 waves persisted with the same propagation speed (Figure 1A). Lack of the dependence of FBP17 wave speed on actin polymerization rate suggests that the waves are not driven by actin polymerization. In addition, lifetime of FBP17 puncta was prolonged (red arrow in Figure 1A). Together those data suggest that the actin likely plays an antagonistic role in the cortical waves of mast cells.

**Figure 1.**
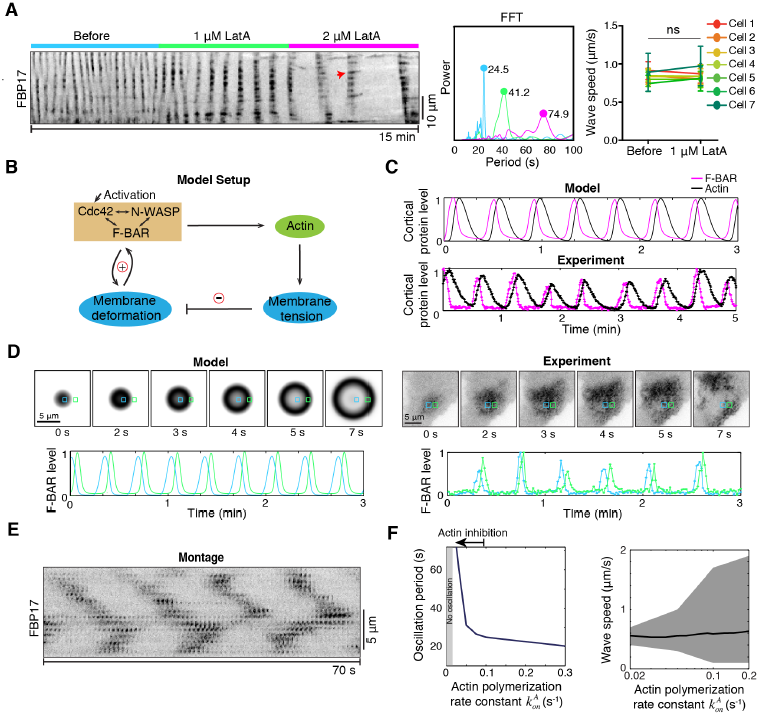
Wave propagation of cortical protein oscillations. (A) Propagation speed of waves is independent of oscillatory periodicity. *Left*, Kymograph of an RBL cell with FBP17-GFP waves treated with titration of Latrunculin A (LatA). Red arrow shows an individual punctum. *Middle*, Fast Fourier transform (FFT) shows the dominant periods (colored dots) of the same cell under each treatment. Colors indicate each treatment as in *Left*. *Right*, Wave speed was not significantly changed after 1 μM LatA treatment. Each pair represents a single cell before and after LatA. Error bar is standard deviation (SD) of wave speed within each cell. (n=4 experiments; ns: no significant difference within each cell; student *t*-test). (B) Model setup of cortical proteins and membrane oscillations. A localized transient pulse of GTP-Cdc42 acts as input. (C) Local cortical protein oscillation. *Top*: The model prediction on the time curves of F-BAR and actin oscillation at the epicenter. *bottom*: Experimental results show similar oscillations of cortical proteins. (D) Cortical oscillation propagation. *Left*: The model predicts a stable phase shift in the cortical oscillation between neighboring areas (the locations marked by □ and □, which are 2.5 μm apart). *Right*: Experimental measurements show stable phase shifts existed between the fixed locations within the propagating wave (the two locations are marked by □ and □, 2.88 μm apart). (E) Montage of FBP17 waves shows both the diffusive population and the immobile puncta. (F) Model results on wave speed and oscillation period. *Left*, Model prediction on dependence of oscillation period on inhibition of actin polymerization. *Right*, Model prediction on insensitivity of average wave speed (black curve) upon variation in actin polymerization rate. There are the maximum and minimum wave speeds for each wave, which together define the range of wave speed (shaded area).

### Mechanochemical Model Recapitulates Wave Propagation

We decided to identify mechanisms that could positively activate wave propagation on the ventral membrane. Considering the facts that F-BAR proteins are members of BAR superfamily proteins that can sense and generate membrane curvature [31–33], and the theoretical concept that membrane shape can feed back to cortical protein recruitment in waves [18,19,21,28,34], we reason that these cortical proteins may mediate membrane shape changes that are important to rhythmic propagation. We set out to determine theoretically whether a model involving membrane shape propagation and curvature-dependent biochemical feedbacks can recapitulate our experimental observations quantitatively.

The mathematical essence of the model is formulated into a set of partial differential equations that govern membrane shape changes, cortical reaction and molecular diffusions (Equations [2.1] - [2.8] and Section 2 of Materials and Methods). Briefly, the model assumes a transient activation signal that increases the local amount of GTP-bound Cdc42 at the cortex (Figure 1B). Active GTP-Cdc42 at the membrane is assumed to recruit effector proteins N-WASP and F-BAR domain-containing proteins (e.g., FBP17, CIP4 or Toca-1) [30,35,36], which we refer to a generic entity as F-BAR. Waves of active Cdc42 were synchronized with the FBP17 and N-WASP waves [30]. A similar activation signal may be evoked by the intrinsic cellular state via a self-assembly process that does not require external stimulation, as in the case of spontaneous travelling waves [30]. Oscillations in the model arise due to the non-linear mechano-chemical pathways downstream of Cdc42 activation.

Cortical Cdc42/N-WASP/F-BAR (CWF) complex could elicit two responses in the model. First, when the F-BAR level exceeded a threshold value, F-BAR would generate membrane shape changes. We postulate that the resulting local curvature induces further F-BAR recruitment to the cortex due to their curvature-sensing ability [37], which serves as the basis of positive feedback between activation of CWF complex and membrane shape changes. Second, active N-WASP recruits Arp2/3 complex and promotes actin polymerization [38–40]. This is based on the observation that waves of active Cdc42 preceded actin waves [30]. Increased actin polymerization decreases local membrane deformability and antagonizes F-BAR cortical recruitment [41–43]. This results in a delayed negative feedback of actin polymerization to cortical recruitment of CWF complex, forming the basis of the temporal oscillations. The model shows that oscillations arise with parameters that are within the range of measured values under physiological conditions (Tables S1 and S2). In agreement with experiments, it successfully recapitulates the local oscillation of cortical F-BAR and actin with phase shift by total internal reflection fluorescence (TIRF) microscopy (Figure 1C).

According to the model, travelling wave appearance of cortical proteins can develop as the out-of-plane membrane undulation conveys the membrane shape deformation in the activation patch to the neighboring cortical area, where such deformation invokes mechanochemical feedback to locally recruit proteins from the cytoplasm (Figure 1D). As such, wave propagates well beyond the original cortical activation site, like a ripple moving in the pond. The model predicts two types of propagation depending on the rate of local cortical activity (Figure S1A). For the “ripple wave”, the wavefront displays saltatory instead of smooth movement. That is, both the speed and the cortical protein level at the wavefront oscillate, and there is a phase shift between them (Figure S1A, *Right*). The saltatory behavior of the wavefront stems from the fine balance between rate of cortical protein recruitment and rate of the attended membrane shape change. Away from the balance arises the “continuous wave”, in which the wavefront travels at a roughly constant speed and cortical protein intensity (Figure S1A, *Middle*). The buildup of cortical protein, which drives nucleation, is in synchrony with wave propagation. If the cortical protein level at the wavefront is below threshold, the wave is predicted to decay at the edge of the cortical patch and eventually die out (Figure S1A, *Left*). Those predictions match with experimental observations (Figure S1B).

The delayed actin-dependent negative feedback supports a travelling wave regime in a wide range of parameter space (Figure S2). While wave propagation has contributions from both lateral diffusions and the local mechanochemical reactions through recruitment from the cytosol, our calculation shows that ~90% of the increase in the local protein concentration at the wave front stems from the recruitment reactions from cytoplasm, while the protein lateral diffusion contributes to ~10% (Figure S3). This suggests that the wave propagation reflects more of local protein assembly from cytosol than protein diffusional drift along the cortex. This prediction is consistent with the lack of laternal movement of the individual FBP17 puncta in the wave (Figure 1E). The model also recapitulates the dependence of oscillation period and the relative insensitivity of propagation speed on actin polymerization rate (Figure 1F), in consistent with LatA experiments (Figure 1A).

### Membrane Shape Oscillations Accompany Traveling Wave Propagation

We next attempted to further validate the model experimentally by testing several predictions. One defining feature is that membrane shape changes are involved in wave propagation. Figure 2A shows membrane shape deformation accompanied the traveling waves over time and space in the model. Cortical protein recruitment was in synchrony with membrane shape change, and marks the wave front propagation (Figure 2B). Note that, the local F-BAR level closely followed that of membrane curvature, but not the membrane height (Figure 2B). To obtain information on the local membrane shape and its relation to the local rhythm of cortical protein recruitments, we simultaneously imaged FBP17 by TIRF microscopy and the distance between cell membrane and the substrate by surface reflection interference contrast (SRIC) microscopy [24]. We found that local membrane shape oscillated and the deformation strongly correlated with the location of the wave marked by cortical FBP17 density (Figure 2C). In addition, both theory and experiment showed a longer duration of membrane height oscillation compared to that of F-BAR oscillations (Figure 2D), due to the delay between changes in membrane curvature and membrane height (Figure 2B). Membrane oscillation had a phase lag relative to F-BAR (Figures 2D, S4A and S4B). Importantly, F-BAR was in the same pace with the rate of membrane height changes (d*h*/d*t*) at any fixed location (Figures 2D and S4C), consistent with the reciprocal interaction of F-BAR recruitment and local membrane curvature underlying the wave propagation. Moreover, actin was anti-correlated with the rate of membrane height changes (d*h*/d*t*) (Figure S4A and S4C), indicating its antagonistic effects on membrane.

**Figure 2.**
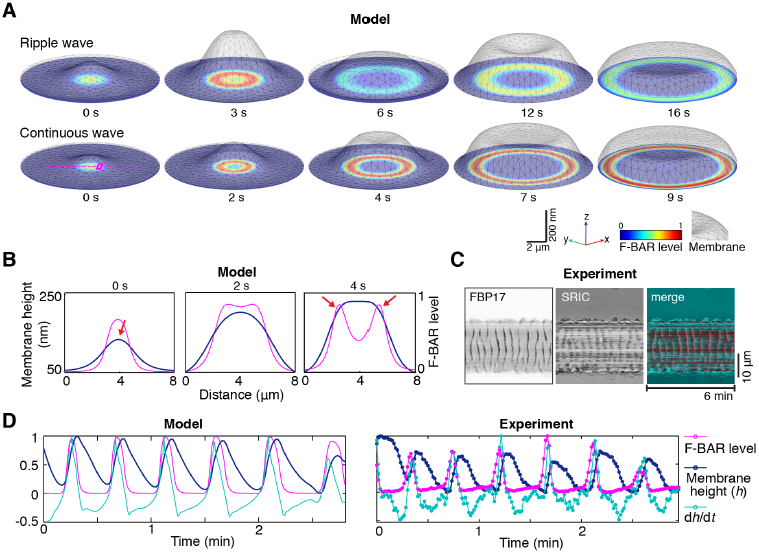
Membrane shape changes accompany traveling wave propagation. (A) Snapshots from model prediction show that the local membrane shapes correlate with the F-BAR cortical density at the wavefront during wave propagation. The grayish meshes represent the membrane shapes in three-dimensional space (X, Y, Z) with a perspective angle. The color maps are the corresponding F-BAR densities on the membrane, which is projected on the two-dimensional x-y plane. The scale bar for the x-y dimension is 2 μm and for the z-dimension is 200 nm. (B) A zoom-in view of the local cortical rhythm along the magenta dash line across the cortex marked in the left panel of Figure 2A at time points t = 0 s, 2 s and 4 s. Blue lines: The local membrane shape. Magenta lines: F-BAR cortical level. The red arrows mark the locations with the largest membrane curvature. (C) Kymographs of the F-BAR cortical waves and the corresponding relative membrane height (SRIC) waves. Scale bar: 10 μm. (D) Local cortical oscillations of F-BAR and membrane. *Left*: Model prediction on the time curves of local F-BAR cortical level, membrane height, and the rate of membrane height changes (d*h*/d*t*) at the fixed location (marked as the magenta square in the left panel of Figure 2A). The membrane height was normalized relative to the highest membrane height. *Right*: Experimental measurement of the spatio-temporal changes in F-BAR cortical level, membrane height (by SRIC), and the rate of membrane height changes (d*h*/d*t*) at a fixed location during traveling wave propagation.

### Curvature Preference of F-BAR is Critical for Wave Propagation

As membrane shape did change detectably along the wave propagation path, we further determined whether membrane shape changes were necessary for wave propagation. In the model, wave propagation requires a threshold level of curvature-dependent cortical protein recruitment, below which it can not sustain the residual membrane shape deformation at the edge of the original cortical patch of activation (Figure 3A). We first examined the effect of removing F-BAR proteins on the formation of traveling waves. We reduced the level of endogenous curvature-generating proteins by shRNA. Knocking down FBP17 or its homolog CIP4 alone did not significantly reduce the percentage of the cells with traveling waves, but double knocking down (DKD) FBP17 and CIP4 abolished the formation of traveling waves (Figure 3B). This indicates some redundancies among the Toca family of F-BAR proteins and demonstrates their essential role as a collective entity of the Cdc42 interacting F-BAR proteins in the traveling waves.

**Figure 3.**
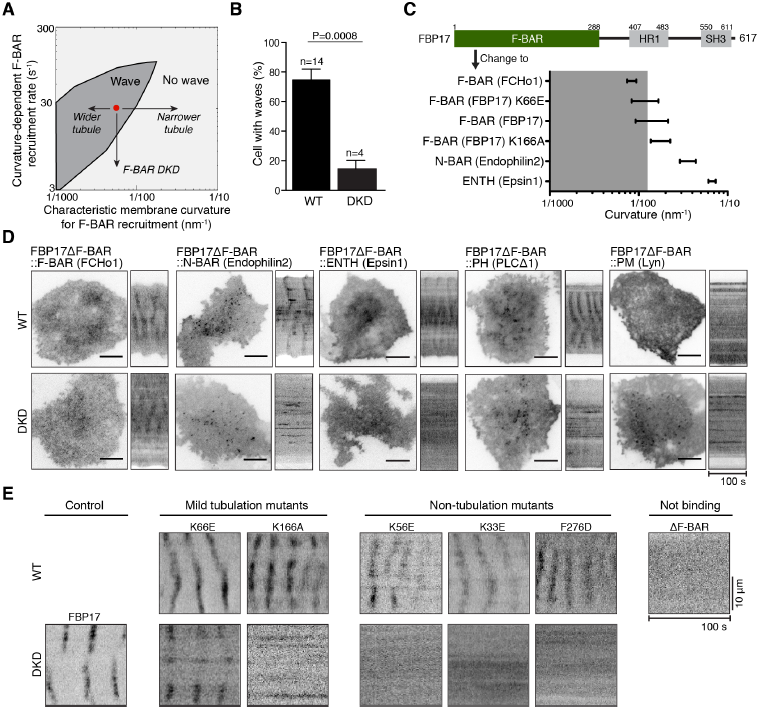
Wave propagation requires F-BAR and its curvature. (A) Model phase diagram of the dependence of traveling waves on a proper balance between the curvature-dependent F-BAR-recruitment rate and the absolute characteristic membrane curvature that recruited F-BAR. The red dot represents the model parameter set that generated the nominal model results in Figure 2. (B) Double knockdown (DKD) of FBP17 and CIP4 by shRNA causes a reduction (*P*=0.0008, student t-test) in the percentage of cells with active Cdc42 (CBD-GFP) waves (4 experiments with 39 cells in total), comparing to wild-type (WT) cells (14 experiments with 133 cells). (C) F-BAR domain of FBP17 is replaced with other membrane-binding motifs. Ranges of curvature (1/diameter) preferences of different membrane-binding domains [51,70–72] are plotted. Grey area indicates the curvature range that allows waves, corresponding to the grey area of Figure 3A. (D) The curvature of F-BAR domain is critical for wave. A micrograph and a kymograph for each mutant protein both in WT and DKD conditions are shown. Only the mutant protein with an F-BAR (FCHo1) domain could rescue wave formation in DKD cells. Scale bar: 10 μm. (E) The ability of point mutants of F-BAR domain of FBP17 to localize to the waves in WT cells or rescue wave formation in DKD cells correlates with their tubulation activities *in vitro*. Representative kymographs of cells expressing point-mutants or full length construct of FBP17-GFP in WT cells and DKD cells are shown. Mutants are separated into four groups based on curvature-generating abilities. Scale bar: 10 μm.

To determine whether the F-BAR domain of FBP17 was necessary or sufficient for the FBP17 wave, we generated multiple versions of truncated FBP17. Deletion of any of the three structural domains (F-BAR, HR1, or SH3) abolished the wave-like appearance of cortical FBP17 in wild-type cells (Figure S5), indicating that the F-BAR domain was necessary, but not sufficient for generating the traveling waves. ΔSH3 and ΔHR1 mutants each formed some stable puncta on cell membrane, but ΔF-BAR did not (Figure S5). Because the SH3 domain and HR1 domain interact with actin machinery, the formation of stable FBP17 puncta in ΔSH3 and ΔHR1 mutants suggests the requirement of a link to actin for dissociation of FBP17 from cortex, thus supporting the actin-mediated negative feedback proposed by our model.

Because F-BAR proteins could function in regulating actin dynamics in addition to their membrane-remodeling ability, the indispensable role of F-BAR proteins in wave propagation did not necessarily prove a requirement for membrane shape changes. To further explore whether the wave propagation requires changes in membrane shape, we generated a series of domain-swapping mutant proteins which differed from FBP17 only in their curvature preferences and tested whether they could functionally replace FBP17 in the waves. Two types of mutants were designed based on biophysical properties of known curvature-generating proteins, which covered a range of curvature preferences from 0.1 nm^-1^ to 0.01 nm^-1^ (Figure 3C). First, we replaced the F-BAR domain of FBP17 with other lipid-binding domains of different shapes: F-BAR domain of FCHo1, BAR domain of Endophilin2, ENTH domain of Epsin1 (Figures 3C and 3D). These domains represented three major classes of curvature-generating domains (F-BAR, BAR and ENTH) [70–72]. Phospholipid-binding but curvature-insensitive PH domain of phospholipase C [73] or membrane-targeting sequence of tyrosine-protein kinase Lyn (Lyn10) were introduced as controls. Among these, the mutant with the constitutively membrane-targeting Lyn10 did not form waves in wild-type (WT) cells, consistent with a requirement of F-BAR dissociation from the membrane for waves (Figure 3D, *top*). The rest of the domain-swapped mutants still localized to the waves in WT cells, indicating that these mutants were properly folded functional proteins. However, in DKD cells, only other F-BAR (F-BAR of FCHo1) could rescue wave formation (Figure 3D, *bottom*). The fact that the BAR (from Endophilin2) or ENTH (from Epsin1) domain-swapped mutant could not rescue wave formation suggests that a specific range of curvature (similar to or lower than that of the F-BAR domains) is required for wave formation.

For the second type of mutants, we introduced single point mutations to perturb the curvature preference of the F-BAR domain of FBP17 (Figures 3C and 3E). Mutants K56E, K33E and F276D had paralyzed tubule-generating ability in living cells [31,51]. Of particular interests are mutants K166A and K66E, which generated narrower (higher curvature) and wider tubules (lower curvature) compared with wild-type F-BAR domain, respectively, in *in vitro* liposome tubulation assays [51]. Both mutants K166A and K66E were localized in waves in WT cells (Figure 3E, *top*), confirming that they were functional. However, only K66E point mutant of FBP17 (of shallower curvature preference) could rescue wave formation in DKD cells (Figures 3E, *bottom*). Together with the domain-swapping data (Figure 3D), these results indicate that the shape of the lipid-binding domain of FBP17 is important for wave formation, regardless of the origin of the curvature preferences being the intrinsic protein shapes or the conformation of protein oligomers. We conclude that membrane shape-mediated feedback is actively involved in the cortical wave propagation.

### Wave Propagation Requires Optimal Plasma Membrane Deformability

The requirement of membrane shape-mediated feedback confirmed the mechanochemical nature of the wave propagation. An independent test for this was on the mechanosensitivity of the waves. As membrane deformability critically determined rhythmic propagation of cortical protein recruitment in our model, the wave propagation was predicted to diminish with increasing or decreasing membrane tension (Figure 4A). Hence, we investigated the effect of membrane mechanics on the waves. We conducted osmolarity shock experiments by cyclically perfusing cells with hypotonic or hypertonic buffers; these treatments increased or decreased membrane tension respectively, which subsequently affected membrane shape deformability. The kymographs show that osmolarity changes reversibly inhibited rhythmic wave propagation (Figure 4B). Osmotic treatment could lead to additional effects including cell volume and cytosolic concentration changes (Figure S6). To circumvent these potential side effects, we used the surfactant deoxycholate (DC) to specifically reduce the plasma membrane tension without changing cell volume [74]. We observed that FBP17, actin, and membrane shape waves disappeared (Figure S7) in a DC dose-dependent manner (Figure 4C). Such effects were reversible on the time scale of seconds upon DC washout, eliminating possibilities of permanent membrane compositional changes. Waves experiencing moderate DC or hyper-osmotic treatment displayed no significant changes in propagation speed, but prolonged oscillation periods (Figure 4D), indicating that the shape-mediated mechano-chemical feedback quantitatively determines cortex excitability.

**Figure 4.**
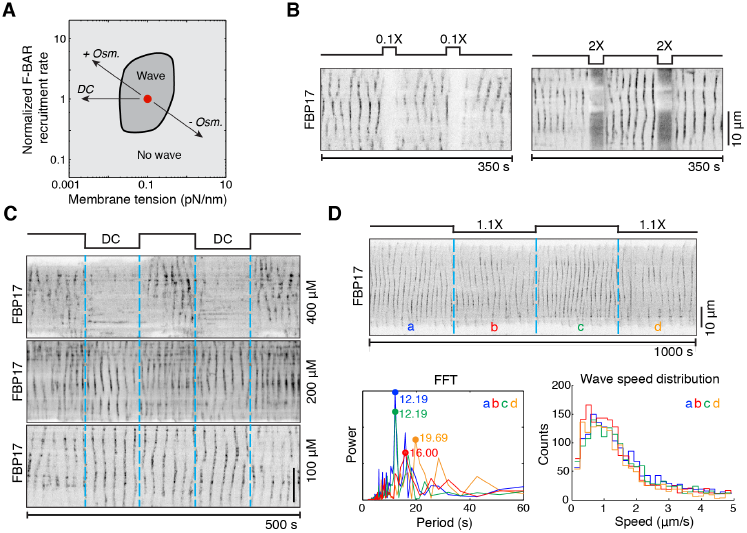
Membrane mechanics is essential for the wave propagation. (A) del phase diagram of the dependence of traveling waves on the membrane tension and the F-BAR cortical recruitment rate. The red dot represents the model parameter set that generated the the nominal model results in Figure 2. (B) Osmotic shock reversibly inhibits traveling waves as shown by the FBP17-GFP kymographs. cells were cyclically perfused with isotonic (300 mOsm) buffer for 100 s and hypotonic (30 mOsm) or hypertonic (600 mOsm) buffer for 25 s. (C) Kymographs of FBP17-GFP shows that surfactant deoxycholate (DC) reversibly inhibited traveling waves in a dose-dependent manner. Cells were cyclically perfused with isotonic buffer without DC for 100 s and isotonic buffer with DC at different concentrations (400 μM, 200 μM, and 100 μM, respectively) for 100 s. (D) Mild osmotic shock changes oscillation period but not wave speed. *Top*: Kymograph of FBP17-GFP waves in cell that was cyclically perfused with mild hyper-osmotic buffer for 250 s each duration. In duration a and c, the cell was perfused with 280 mOsm buffer, while in duration b and d, cells were perfused with 310 mOsm buffer. *Bottom left*: FFT shows the dominant periods in each duration. *Bottom right*: Wave speed distribution in each duration. Colors of plots indicate durations as in *Top*. For (B)–(D), All the schematics above kymographs illustrate putative changes in cell membrane tension. Scale bar: 10 μm.

## Discussion

Taken together, our results present a biological scenario where membrane shape-mediated mechano-chemical feedback is critical for spatiotemporal pattern formation in the dynamical systems. Our data agree with the sequential actin assembly model proposed for actin waves in *Dictyostelium* [10] and *Xenopus* [75], which is in contrast to collective pattern formation of actin that requires bulk actin flow or transport of actin filaments [76]. Waves are not driven by the protrusive forces of actin on the membrane, as suggested by previous models [8,28,77]. The antagonistic role of actin on cortical wave is similar to what was proposed in other systems [4,12,75,78], but the activating signal in those models were assumed to be purely chemical. Here both our model and experiments show that the plasma membrane acts as an active medium for cortical traveling waves. Membrane shape changes establish a spatial cue for cortical protein recruitment, which reciprocally governs the cortical protein dynamics, rendering the plasma membrane an excitable system. Changes in membrane shape and consequently wave propagation depend sensitively on both the dynamic protein membrane interaction and membrane deformability. Softer membrane confers a faster F-BAR-mediated membrane shape deformation that reciprocally speeds up F-BAR recruitment to the cortex, which ends up deforming the membrane in a highly localized manner. Because membrane tension is low, smoothing of this deformation is slow. This introduces a further delay in the actin polymerization-mediated negative feedback on F-BAR, prolonging the oscillation period. However, if the membrane tension is reduced even further, it will not be able to smooth out the membrane deformation on a relevant timescale and consequently the cortical rhythm is disrupted. In this latter case, F-BAR
 puncta remains non-oscillatory and non-propagating, which phenocopies the non-propagating puncta induced by the BAR domain mutants. Conversely, stiffening membrane (as in hypotonic treatment) inhibits membrane shape changes and F-BAR-protein recruitment. When the F-BAR level is lower than the threshold value, cortical oscillation stops, and wave decays.

Travelling wave has been proposed to correlate information processing at integrated level in giant single cells or multi-cellular systems [79–81]. Recently, travelling waves of actin and GTPase including Rho [75], Rac [82] and Cdc42 [83] have emerged as hallmarks and likely regulators for cell cycle regulation, cell migration and chemotaxis. Mechanochemical sensitivity of F-BAR proteins also appears to be important for cell polarity [84]. Such curvature-mediated spatial coupling, manifested as travelling waves, theoretically allows cell to rapidly scan its surface for mechanical and chemical cues. This provides an efficient mechanism of cortical signal transmission that can be widely applicable in diverse cellular contexts.

## Materials and Methods

### Section 1. Experimental methods and data analysis

#### Cell culture and transfection

RBL-2H3 cells (tumor mast cells) were maintained in monolayer in MEM medium (Life Technologies, Carlsbad, CA) containing 20% FBS (Sigma-Aldrich, St. Louis, MO) and 50 μg /mL gentamicin (Life Technologies). Cells are harvested with TrypLE™ Express (Life Technologies, Carlsbad, CA) 2-5 days after passage. For transient transfections, electroporation with Neon transfection system (Life Technologies, Carlsbad, CA) was used. After transfection, cells were plated at subconfluent densities in 35-mm glass bottom dishes (MatTek, Ashland, MA) or on round coverslips in a 12-well plate overnight. Before imaging, cells were washed twice with Tyrodes buffer (135 mM NaCl, 5.0 mM KCl, 1.8 mM CaCl_2_, 1.0 mM MgCl_2_, 5.6 mM glucose, 20 mM Hepes (pH 7.4)). For double knock-down experiments, cells transfected with four HuSH-29 shRNA (Origene, Rockville, MD) at 0.05 μg/μl for each shRNA targeting rat CIP4 (TATGCGAAGCAACTCAGGAGTCTGGTGAA, GCCGCAGAGTCCGTGGATGCTAAGAACGA) and FBP17 (ATGGACGCCGACATCAATGTGACCAAGGC, TGAAACGCACGGTGTCAGACAACAGCCTT) were incubated for 48 hours before imaging. In all experiments, cells were sensitized with mouse monoclonal anti-2,4-dinitrophenyl IgE (Sigma-Aldrich, St. Louis, MO) at 0.5 μg/ml overnight and stimulated with 80 ng/ml multivalent antigen, 2,4-dinitrophenylated bovine serum albumin (DNP-BSA, Life Technologies, CA) before experiments. For experiments using inhibitors, latrunculin A (Sigma-Aldrich, St. Louis, MO) was diluted from the stock and added to the cells at a final concentration of 1 or 2 μM.

#### Molecular Cloning and plasmids

Domain-swapping mutants of FBP17 were generated by replacing F-BAR domain (8-288 amino acids, a.a.) of human FBP17-GFP with F-BAR domain (1-275 a.a.) of mouse FCHo1, BAR domain (1-241 a.a.) of rat Endophilin2, ENTH domain (1-144 a.a.) of rat Epsin1, PH domain (1-170 a.a.) of human PLCΔ1, membrane targeting sequence (1-10 a.a.) of human tyrosine-protein kinase Lyn, respectively. DNA sequences corresponding to 1-7 a.a. and 289-295 a.a. of FBP17 were kept as homologues regions for overlapping PCR. Point mutations on F-BAR of FBP17 were generated using Phusion Site-Directed Mutagenesis Kit (Thermo Scientific, Waltham, MA). Constructs for the following proteins were kind gifts: Lifeact-mRuby from Dr. Roland Wedlich-Soldner (Max Planck Institute of Biochemistry, Martinsried, Germany); FBP17-GFP, mCherry-FBP17, CBD-GFP (a Cdc42 activity probe) and mCherry-actin were from Dr. Pietro De Camilli (Yale University School of Medicine, New Haven, CT) as previously described [30]. All plasmids were sequenced to vindicate their integrity.

#### Total internal reflection fluorescence (TIRF) microscopy

A Nikon Ti-E inverted microscope equipped with perfect focus system, iLAS2 motorized TIRF illuminator (Roper Scientific, Evry Cedex, France) and an Evolve 512 EMCCD camera (Photometrics, Tucson, AZ) was used. All images were acquired through a TIRF-CFI objective (Apochromat TIRF 100XH N.A. 1.49, Nikon). Samples were excited by 491 nm (100 mW) or 561 nm (100 mW) laser, reflected from a quad-bandpass dichroic mirror (Di01-R405/488/561/635, Semrock, Rochester, NY). The emitted light was acquired after passing through an emission filter (FF01-525/45 for GFP or FF01-609/54 for RFP/mCherry, Semrock, Rochester, NY) located on a Ludi emission filter wheel. MetaMorph 7.8 software (Molecular Device, Sunnyvale, CA) was used for image acquisition. Samples were maintained at 37 °C throughout the experiments using an on-stage incubator system (Live Cell Instrument, Seoul, South Korea).

#### Surface reflective interference contrast (SRIC) microscopy

Same setup as TIRF was used except that a mercury lamp (X-Cite 200DC, 200 W, Excelitas, Waltham, MA) was used as the light source. The mercury lamp had a spectrum output at wavelengths between 340 nm to 800 nm. A bandpass filter (609/54; Semrock, Rochester, NY) and neutral density filters (ND4) were used. Near-parallel rays produced by closing aperture diaphragm obliquely illuminated the sample through the TIRF-CFI objective (100X N.A. 1.49, Nikon, Japan). Reflected light rays from two interfaces (between coverslip and solution and between solution and plasma membrane) interfered with each other, creating a bright or dark patch when they were constructive or destructive, respectively. In order to perform sequential TIRF/SRIC with fast frame rate, mechanical changes of the optical parts were reduced to a minimum level. A 30/70 beam splitter was used at the back port of the L-shape fluo-illuminator of Nikon microscope to combine the white light source (mercury lamp used for SRIC) and the lasers (used for TIRF). The same quad-bandpass dichroic cube (Di01-R405/488/561/635, Semrock, Rochester, NY) used for TIRF was used for SRIC, which allows an estimated 2% of incident light (609/54) to be reflected during the SRIC acquisition. Similar results were obtained using a Nikon SRIC cube composed of excitation filter 535/50, dichroic mirror DM400 and neutral density filter ND16.

#### Perfusion experiments

Transfected cells were seeded on 20 mm round coverslips in a 12-well plate overnight. Prior to imaging, the coverslip was transferred to a custom perfusion chamber (Chamlide, Live Cell Instrument (LCI), Seoul, South Korea) placed on the heated stage of a Nikon Ti-E inverted microscope. A multi-valve perfusion control system (MPS-8, Live Cell Instrument (LCI), Seoul, South Korea) was used to switch rapidly between solutions flowing into the chamber and over the cells. The perfusion system was connected to a computer and controlled by MetaMorph software (Molecular Device, Sunnyvale, CA) in synchronization with image acquisition. A typical flow rate was ~ 0.3 mL/min. For the inner channel dimension of 10 mm (L) × 2 mm (W) × 0.2 mm (H), the flow rate was calculated to be 12.5 mm/s in the channel, which means the solution in the chamber was fully exchanged in less than one second. The following solutions were used: Tyrodes buffer, Tyrodes buffer supplemented with 400 μM deoxycholate (DC), hyper-osmotic Tyrodes buffer with 2X solute concentration and hypo-osmotic Tyrodes buffer diluted with distilled deionized water to 0.1X/0.5X concentration. pH of all buffers was adjusted to 7.4. Osmolarity was confirmed using an osmometer (Osmomat 030, GONOTEC GmbH, Berlin, Germany).

#### Image analysis

An ImageJ-based software Fiji [85] was used to generate kymographs, montages and Z-stack projections. MATLAB (MathWorks, Natick, MA) was used for data quantification and plotting. To identify frequency information for time-series intensity profile of ROIs of cells, the built-in fast Fourier transform function in MATLAB was used. We determined the instantaneous wave speed by the following procedures. To avoid complications arising from the interactions between multiple waves, we only chose the well-defined single waves to determine the wave speed. We segmented individual waves by setting the intensity threshold level just above the average background intensity. Next, wave centroid for Figure 4D (or the most advanced point of wavefront patch for Figure 1D) ***r***_*c*_(*t*_*i*_) of the segmented region for each time frame *t*_*i*_ was calculated. From the trajectory of the wave centroids over time, we obtained the instantaneous wave propagation direction, and wave speed, as determined by *v=*|***r***_*c*_*(t*_*i+1*_*)-****r***_*c*_*(t*_*i*_)|/*(t*_*i+1*_*-t*_*i*_).

Moreover, the robustness of our wave tracking results depended on the chosen segmentation threshold value. To ensure that the tracking can give consistent results, we varied segmentation threshold values to determine the optimal range. Our calculation showed that at each time point of a given wave, the instantaneous wave speed from our tracking analysis indeed varied with chosen segmentation threshold values. We then used clustering analysis to determine the optimal segmentation threshold range, within which the speed variation was minimal and, thus, our tracking results for this individual measurement converged. By overlapping the individual range from every wave at every time point of the same cell, the overall optimal threshold range for this cell was determined. This optimal range could vary in different cells in part due to the different background intensities. Throughout the tracking, we thus chose a fixed segmentation threshold value within the optimal range that is independently determined for individual cells.

### Section 2. Model equations and parameters

The model describes rhythmic propagation emerging from the interplay between membrane-bound chemical reactions and membrane shape changes. For cortical protein recruitment from and turnover into the cytoplasm, the cytoplasm is treated as an unlimited reservoir. Once recruited to the ventral cortex, these proteins diffuse laterally; notably, cortical diffusion is much slower than cytoplasmic diffusion [56,86]. The model assumes the membrane as an elastic sheet. As the model concerns the cortical dynamics occurring on the ventral membrane, membrane shape is governed by membrane tension, bending modulus, membrane-substrate adhesion, and mechanical action of membrane-bound proteins, including F-BAR-mediated membrane shape deformation and actin-mediated enhancement of membrane tension. The resulting membrane shape promotes F-BAR recruitment, which is depicted with Michaelis-Menten-like kinetics.

We determine the dynamical evolution of the system by integrating these equations from an initial condition: a flat membrane patch (40 μm across) and no cortical proteins. In the simulation, this membrane patch represents the ventral side of the plasma membrane, beyond which the model imposes a pinning boundary condition that ramps up a large resistance force, restoring the membrane height back to baseline, reflecting the hindrance effect of the cell edge, as the traveling wave in our system is observed on the cell ventral side. With an input of a localized transient pulse of GTP-Cdc42, the model output at each time step is membrane shape and local protein densities.

In the following, we describe the formulation of model equations. We focus on the simplest mechanism and only include the most essential components suggested by experiments. These components are: Cdc42, N-WASP, F-BAR, actin, Arp2/3, and membrane shape. Below, we will translate this qualitative picture into a set of coupled partial differential equations (PDEs). All the chemical reactions are modeled by Michaelis-Menten type kinetics, in which the local membrane shape influences the on rate of F-BAR. Conversely, the local F-BAR level sculptures the shape of the local membrane, and the local actin level modulates the membrane tension. In this way, the membrane mechanics is coupled with the local chemical reactions. These PDEs thus depicts the spatio-temporal dynamics of the mechanochemical coupling of the membrane shape and the membrane-bound chemical reactions. The model parameters are listed in Tables S1 and S2.

*Cdc42 dynamics:*

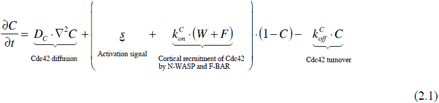

*N-WASP dynamics:*

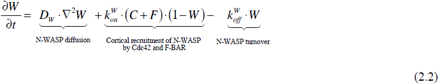

*F-BAR dynamics:*

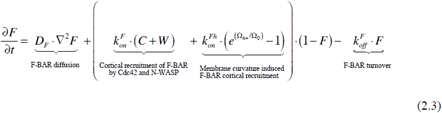

Here, the term 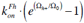 depicts the membrane curvature-dependence of F-BAR cortical recruitment [49,50,57,87]. Ω_0_ is the characteristic membrane curvature, beyond which F-BAR cortical recruitment is significantly increased. We would like to emphasize that the F-BAR in the model refers to a collective functional entity of multiple F-BAR domain proteins, instead of referring to any individual F-BAR protein. This is consistent with our double knockdown experiment results in Figure 3B. Consequently, the characteristic membrane curvature Ω_0_ is within the range of the values measured/inferred from different F-BAR proteins. Because F-BAR is more flexible than BAR-domain proteins, it could bind to the membranes with different curvatures that correspond to vesicle sizes ranging from 60 nm to microns in diameter [31,49,50]. An *in vitro* experiment demonstrated this curvature sensing of FCHo (a F-BAR protein), which corresponded to the size of lipid vesicle ~ 200-400 nm in diameter [49,50]. In the model, we thus chose the nominal value of Ω_0_ to be ~ 1/200 nm^-1^. In addition, our phase diagram study suggests that the model essence persists over a broad range of this threshold curvature Ω_0_ (Figure 3A).

*Arp2/3 dynamics:*

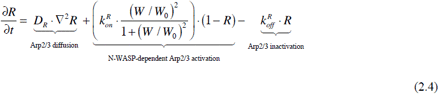

Note that N-WASP is known to activate Arp2/3 complex as a dimer [40]. Therefore, the Hill coefficient is 2.

*Actin dynamics:*

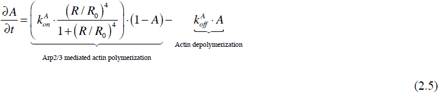

While many proteins are at play, the equation (2.5) with Hill coefficient = 4 is the simplest form that is meant to capture this nonlinearity. Any cooperativity with Hill coefficient > 2 would preserve the model essence on the traveling waves (data not shown). We note that Arp2/3-mediated branched actin polymerization is autocatalytic. It involves nucleation phase with latency followed by rapid actin polymerization and, hence, is highly non-linear [38]. The combination of equations (2.2), (2.4) and (2.5) phenomenologically describes the nonlinear dynamics of actin polymerization with an effective time delay that lags behind the N-WASP activation.

*Membrane shape dynamics:*

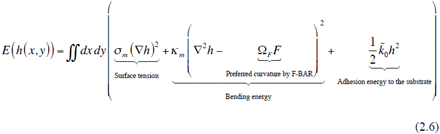

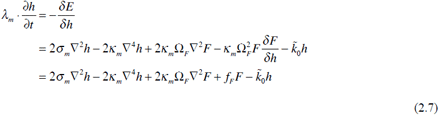

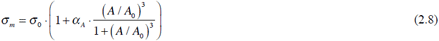

Equation (2.6) describes the Helfrich membrane free energy *E*, where *h* is the membrane height in the z-direction under Monge Gauge, representing the membrane shape over time and space. ∇^2^*h* is the mean curvature, and the preferred curvature by F-BAR is linearly proportional to the local F-BAR level. *k*_*m*_ is the bending modulus and *σ*_*m*_ is the surface tension. 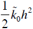 is the adhesion energy between the membrane and the substrate, given that the traveling waves under study are on the ventral side of the cell membrane.

Equation (2.7) is the Langevin equation. It stems from the functional derivative of the free energy functional with regard to the variation of the membrane height *h*, which characterizes the dynamics of membrane shape change. *λ*_*m*_ is effective viscous drag coefficient for membrane shape changes; it combines the membrane resistance and fluid drag from outside, whose value is measured to be ~ 2×10^9^ *Pa*.*s/m* [48]. In addition, the functional derivative 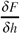 at the membrane equals to the derivative of the F-BAR concentration field in the normal direction of the membrane surface. The F-BAR concentration in this direction has a gradient low in cytoplasm and high at the plasma membrane, so it is negative. In the simplest approximation, we assume that this gradient at the membrane is a constant -*g*_F_ with *g*_F_ >0. Combined together, 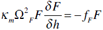.

Here, 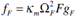, it is essentially the free energy change arising from F-BAR binding to membrane (similar to chemical potential), from which the value of *f*_F_ can be estimated. Furthermore, the F-BAR expression level under this study is low; otherwise, significant membrane tubulation would be evident. In the low F-BAR level limit, the linear term in F always dominates over ∇^2^ *F* term in equation (2.7). In our calculation, we thus ignore the ∇^2^2 *F* term.

Equation (2.8) characterizes the modulation of membrane tension by local cortical actin level. Actin binding to membrane is a complex process and supposed to be highly cooperative. For example, experiments in *Dictyostelium discoideum* showed that actin bound to membrane with Hill coefficient = 3 [88]. We therefore use the hill coefficient = 3 in our model. The qualitative feature of our model remains the same for any hill coefficient > 2 (data not shown). It also has been measured that cortical actin polymerization could increase the measured plasma membrane tension by 2-10 folds [42,43,68,69], which sets the range of the *α*_A_ in equation (2.8).

### Computational scheme

In the simulation, a free boundary condition was imposed. The initial condition was: all chemical components were at the equilibrium; membrane was at the resting state. At time zero, an activation signal was imposed to the system. We solved the coupled PDEs by the finite element solver COMSOL® Multiphysics software (version 4.4, Sweden). The spatial resolution was on the order of 300 nm. All the related parameters were rescaled according to this setup. The time dependent solver was used with relative tolerance of 1×10^-4^. The backward differentiation formula or BDF method was used for time stepping provided by default Multiphysics module of COMSOL. The solver used was MUMPS (multifrontal massively parallel sparse direct solver) from default Multiphysics module of COMSOL.

## Author Contributions

M.S and M.W designed experiments; W.Z and J.L developed the model; M.S and C.T performed the experiments; all analyzed the data and M.S, W.Z, M.W and J.L wrote the manuscript.

## Acknowledgments

We would like to thank Mike Sheetz for suggesting the deoxycholate experiment, Yang Yang, Ding Xiong for sharing experimental results, Emma Feng Yu, Larry Cheung for technical assistance.

---

**Table S1.** Parameters in mechanics

| Parameter Symbol | Parameter description | Measured or estimated value | Value used in the model | Reference |
| --- | --- | --- | --- | --- |
| $\kappa_m$ | Membrane bending modulus | $4.2 \times 10^{-19} J$ | $4.2 \times 10^{-19} J$ | [44,45] |
| $\sigma_0$ | Surface tension | $\sim 10^{-4} - 10^{-5} J/m^2$ | $1 \times 10^{-4} J/m^2$ | [41,44–47] |
| $\lambda_m$ | Membrane viscous drag coefficient | $\sim 2 \times 10^9 Pa s/m$ | $\sim 2 \times 10^9 Pa s/m$ | [48] |
| $f_F$ | Maximum stress exerted by F-BAR | | 750 Pa | Estimated |
| $\tilde{k}_0$ | Membrane-substrate adhesion stress | $\sim 0.05 - 0.25 nN/\mu m^3$ | 0.25 nN/$\mu m^3$ | [44] |
| $\Omega_0$ | Characteristic membrane curvature that recruits F-BAR | 1/50– 1/200 nm<sup>-1</sup> | 1/200 nm<sup>-1</sup> | [31,49–51] |

**Table S2.** Parameters in chemical reactions

| Parameter Symbol | Parameter description | Measured or estimated value | Value used in the model | Reference |
| --- | --- | --- | --- | --- |
| $D_C$ | Diffusion constant for Cdc42 | $\sim 0.02 - 0.06 \mu m^2/s$ | $0.05 \mu m^2/s$ | [52–55] |
| $D_W$ | Diffusion constant for WASP family proteins | $\sim 0.35 - 0.45 \mu m^2/s$ | $0.4 \mu m^2/s$ | [56] |
| $D_F$ | Diffusion constant for F-BAR | $\sim 0.13 - 0.63 \mu m^2/s$ | $0.3 \mu m^2/s$ | [57,58] |
| $D_R$ | Diffusion constant for Arp2/3 | $\sim 0.58 - 0.68 \mu m^2/s$ | $0.6 \mu m^2/s$ | [56] |
| $k_{on}^C$ | Activation rate constant of Cdc42 | $\sim 0.05-5/s$ | 2.5/s | [59] |
| $k_{off}^C$ | De-activation rate constant of Cdc42 | $\sim 0.05-5/s$ | 3/s | [59] |
| $k_{on}^W$ | Activation rate constant of N-WASP | 0.1-3/s | 2.5/s | [60] |
| $k_{off}^W$ | De-activation rate constant of N-WASP | 0.1-3/s | 3/s | [60] |
| $k_{on}^F$ | Recruitment rate constant of F-BAR | | 0.3/s | Estimated |
| $k_{on}^{Fh}$ | Curvature-dependent recruitment rate of F-BAR | | 30/s | Estimated |
| $k_{off}^F$ | Turnover rate constant of F-BAR into cytoplasm | $\sim 0.5-2/s$ | 3/s | [57] |
| $k_{on}^R$ | Recruitment rate constant of Arp2/3 | 0.1-1/s | 0.6/s | [60,61] |
| $W_0$ | Normalized threshold N-WASP level to promote Arp2/3 activation | 0-1 | 0.7 | [60,62] |
| $k_{off}^R$ | De-activation rate constant of Arp2/3 | 0.1-1/s | 0.2/s | [60,61] |
| $k_{on}^A$ | Actin polymerization rate | 0.05-0.25/s | 0.1/s | [25,63–65] |
| $R_0$ | Characteristic Arp2/3 activity to promote actin polymerization | 0-1 | 0.25 | [60,61,66,67] |
| $k_{off}^A$ | Actin depolymerization rate | 0.05-0.25/s | 0.1/s | [25,63–65] |
| $\alpha_A$ | Maximum fold increase in actin-mediated membrane tension | 2-10 | 5 | [68,69] |
| $A_0$ | Threshold actin level to modulated membrane tension | 0-1 | 0.2 | [42,43] |

